# Non-transcriptional gibberellin DELLA signalling for microtubule dynamics

**DOI:** 10.1101/2025.11.11.687800

**Authors:** Yuliya Salanenka, Sravankumar Thula, Alexander Johnson, Han Tang, Lukas Hoermayer, Jiří Friml

## Abstract

Plant cell growth is associated with changes in the dynamics and orientation of the microtubule (MT) cytoskeleton. The plant hormone Gibberellic acid (GA), a central regulator of growth also has an impact on MT arrangements, however the causal relationship between these processes and underlying molecular mechanism remain unclear. Here, examining the *Arabidopsis thaliana* root elongation zone, we show that MT reorientation occurs as a consequence of GA-induced cell elongation and that GA has also a direct effect on the rate of MT polymerization via a non-transcriptional mechanism. Preventing cell elongation blocked GA-induced MT reorganization, whereas GA still promoted elongation in the absence of intact MTs. On the other hand, GA promoted MT dynamics as revealed by the recovery of MTs following depolymerization or photobleaching. Live imaging revealed that GA directly promotes MT polymerization at their plus ends. This GA effect persists even when transcription or translation are inhibited, but still requires the DELLA canonical components of GA signalling and their interactors, the prefoldins (PFDs). Our results support a model where GA-induced DELLA degradation releases ‘in nucleus’ sequestrated PFDs, enhancing their cytoplasmic function to promote MT polymerization. Hence, GA signalling branches into two mechanistically distinct modules: (i) canonical transcriptional pathway that promotes growth and indirectly drives MT reorientation; and (ii) the non-transcriptional DELLA-PFD axis that directly promotes MT polymerization. Such a dual mechanism enables plants to synchronize growth with cytoskeletal remodelling by coupling rapid, cytoplasmic responses to slower transcriptional programs.

## Introduction

Plant growth and development depends on cell expansion and its regulation during various environmental conditions. MTs, mainly the cortical MTs (CMTs) are major structural elements of the cytoskeleton that take part in intracellular transport, mitotic cell division and regulation of cell shape and expansion (Baskin, 1994; Bellinger et al., 2023; Gu et al., 2008; Hsiao & Huang, 2023; Yan et al., 2023). CMTs also guide cellulose synthase complexes, dictating the orientation of cell wall deposition and thereby controlling anisotropic growth (Paredez, Somerville, & Ehrhardt, 2006). The ability of MTs to rapidly reorient in response to mechanical, developmental and environmental cues enables tissues such as the root elongation zone in *Arabidopsis thaliana* to maintain growth flexibility under fluctuating conditions (Hoermayer et al., 2024; Muratov & Baulin, 2015; Yan et al., 2023). CMTs, therefore, act not only as structural guides for cell wall synthesis, but also as signalling compounds that respond to mechanical stress and adapt to growth accordingly (Hamant et al., 2019; Hejnowicz, Rusin, & Rusin, 2000).

Plant hormones are among endogenous regulators of CMT organization. Auxin modulates CMTs’ orientation largely via regulation of growth (Adamowski, Li, & Friml, 2019; Chen et al., 2014). Brassinosteroids (BRs) promote transverse CMT arrays (Delesalle, Vert, & Fujita, 2024), ethylene alters MT stability during differential growth, such as apical hook formation (Le et al., 2005; Polko et al., 2012) and cytokinins influence turnover and orientation of CMTs (Adrian et al., 2023; Montesinos et al., 2020). Gibberellic acid (GA) is also a prominent regulator of growth and other processes including seed germination and flowering (Davière & Achard, 2013; Sun, 2010). It is well established that GA effect on growth correlates with reorganization of CMTs (Baluška, Parker, & Barlow, 1993; Shibaoka, 1974; Wenzel & Wasteneys, 2000). In addition, early reports in monocots suggest that GA stabilizes MTs against depolymerization (Hamada, Mita, & Shibaoka, 1994; Huang & Lloyd, 1999; Mita & Katsumi, 1986; Mita & Shibaoka, 1984). Two major questions arise in this context: (i) What is the causal relationship between GA-regulated cell elongation and CMT reorientation; and (ii) does GA exert direct, growth-independent control over MT dynamics, and what molecular mechanism underlies this effect?

The canonical GA signalling is well known: GA perception leads to degradation of DELLA transcriptional repressors, activating growth-associated transcriptional programs (Davière, De Lucas, & Prat, 2008; Sun, 2010; Ubeda-Tomás et al., 2008; Ueguchi-Tanaka et al., 2005). Notably, DELLAs have been shown to interact with prefoldins (PFDs) (Locascio, Blázquez, & Alabadí, 2013), molecular chaperones also required for tubulin folding and MT organization (Geissler, Siegers, & Schiebel, 1998; Vainberg et al., 1998). The interaction with DELLAs would sequester PFDs in the nucleus and GA-induced DELLA degradation would release them, restoring their cytoplasmic activity (Locascio, Blázquez, & Alabadí, 2013). This provides a possible mode of action on how GA could regulate MTs.

In this study we used the *Arabidopsis thaliana* root elongation zone to dissect how GA influences MT orientation and dynamics. Combining pharmacological, genetic and live cell imaging analysis, revealed that CMT reorientation is a consequence of GA-induced cell elongation. In parallel, GA stimulated MT polymerization at their plus ends through a non-transcriptional, DELLA-PFD-dependent mechanism. These findings establish GA as a dual regulator coordinating transcriptional and cytoplasmic signalling modules to regulate both growth and cytoskeletal remodelling.

## Results

### Cortical microtubules are rearranged as a result of GA-mediated cell expansion

To investigate how growth and MT arrangement are coordinated in response to GA, we examined the dynamics of both CMTs and cell expansion in parallel. We measured the epidermis cell elongation rate in the fast-growing region of roots, both in GA-depleted (pretreatment with the GA biosynthesis inhibitor, Paclobutrazol (PAC) (Rademacher, 1989) and GA-supplemented (GA treatment on PAC-pretreated roots) conditions.

GA-depleted cells showed a strong reduction of cell elongation (20% after 1 hour) and transverse CMTs were mostly absent. The proportion of cells with transverse CMTs increased from ∼20% to ∼100% within the first hour of GA treatment. At the same time, cells elongated by ∼85% on average (Figures 1A, 1B and 1G). These results confirm a correlation between cell expansion and CMT orientation during the root elongation in response to GA, which is consistent with previous studies in different tissues and species (Lloyd et al., 1996; Mita & Katsumi, 1986; Shibaoka, 1974, 1994; Vineyard et al., 2013; Wenzel & Wasteneys, 2000)

**Fig. 1.**
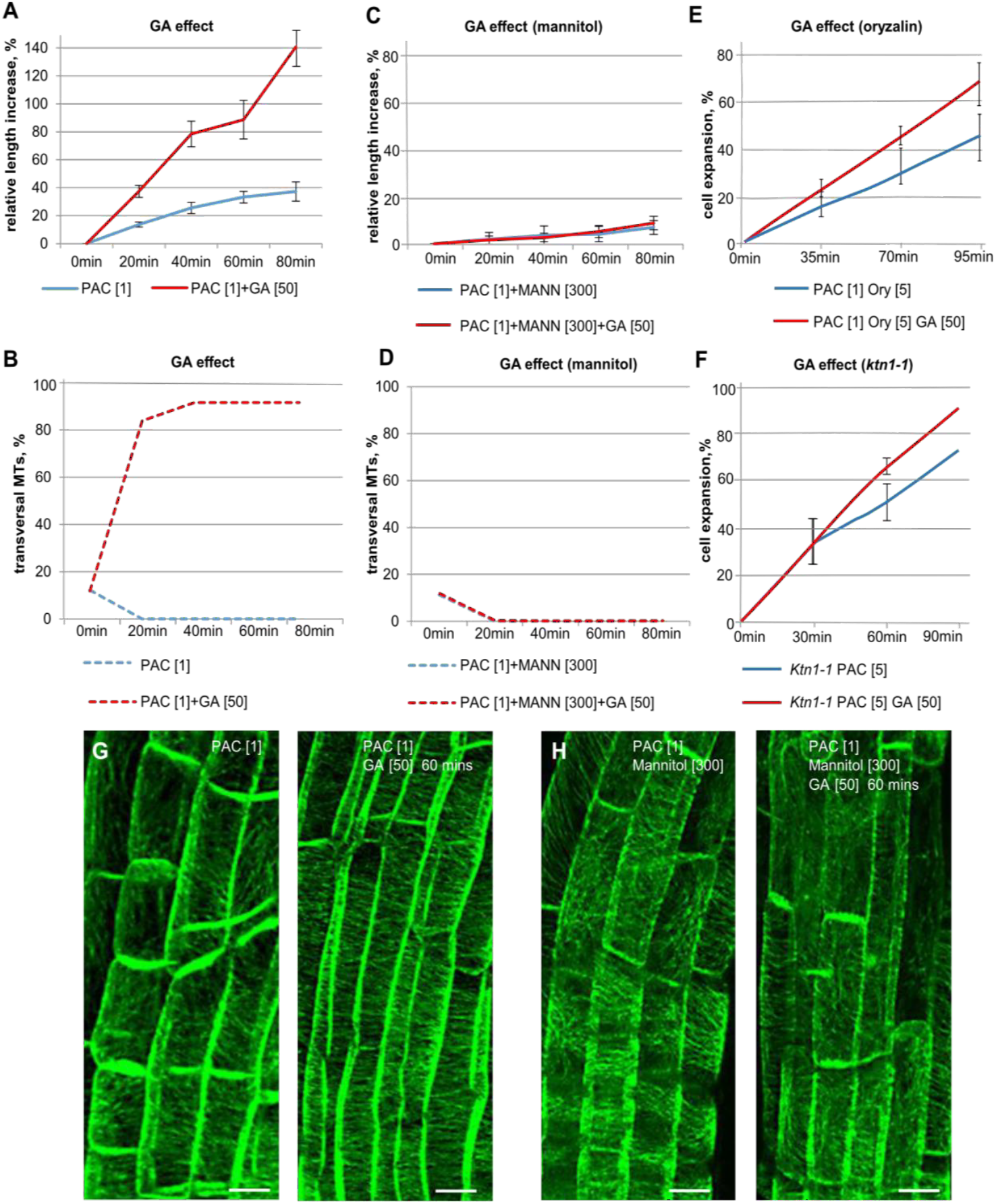
Cortical microtubules rearrangement occurs downstream of GA-mediated cell expansion. **(A**,**B**) GA effect on cell expansion (A) and CMT orientation (B) in epidermis of root elongation zone. Transgenic line carrying MAP4-GFP was pre-treated with 1 μM PAC for 30 mins followed by 50 μM GA treatment as indicated. The percentage of cell expansion in all observed elongated epidermal cells was calculated by dividing the cell length by its original cell length. The percentage of cells with transversal MT reorientation and the cell expansion rate were quantified as described in methods. (**C-F**) GA effects in conditions of inhibited cell expansion by mannitol (300 μM) treatment (C, D) or defective CMTs (oryzalin treatment, E; *ktn1* mutant, F). Roots were pre-treated with PAC and mannitol or oryzalin for 30 mins. In the presence and absence of GA, cell expansion and MT orientation were quantified as described in (A). (**G, H**) Representative images showing the CMTs in indicated conditions, corresponding to (A) to (D).

To investigate the causal link between GA-induced cell expansion and transversal CMT realignment, we perturbed either cell expansion or MT functionality. First, we inhibited cell expansion by mannitol-induced osmotic stress (Kroeger, Zerzour, & Geitmann, 2011; Pritchard, Jones, & Tomos, 1991) (Figure 1C). During the mannitol treatment, there was no rearrangement of CMTs to transversal orientation following GA application (Figure 1D and 1H). This suggests that CMT realignment in response to GA requires cell expansion. In a complementary experiment, we depolymerized MTs using oryzalin treatment (Morejohn et al., 1987) and analyzed the rate of cell expansion with and without GA application. Notably, even after the complete depolymerization of MTs, GA was still able to promote cell expansion (Figure 1E). Furthermore, the *ktn1* mutant, which is unable to establish the proper CMT arrays (McNally & Vale, 1993), still showed increased cell expansion following GA treatment (Figure 1F). Thus, the GA effect on cell expansion can be uncoupled from CMT reorientation.

Overall, these results show that GA primarily promotes cell expansion, and the CMT transversal rearrangement occurs largely as a consequence of this GA-mediated cell expansion.

### GA promotes microtubule network recovery and stability

Previous studies in monocot species, such as onion or maize, have shown that GA can protect MTs from disruption by MT-depolymerizing compounds or by low temperature suggesting that GA has a positive effect on MT stability (Hamada, Mita, & Shibaoka, 1994; Huang & Lloyd, 1999; Mita & Shibaoka, 1984). To further investigate possible GA effects on MT orientation during root growth, we tested whether GA has effect on CMT stability or dynamics.

First, we depolymerized MTs using oryzalin and monitored the rate of MT network recovery in GA-deficient and GA-optimal conditions by following the MT-associated protein, GFP-MAP4 (Marc et al., 1998; Nogales, 2001). In the GA-optimal (mock-treated) roots, newly assembled MTs appeared about 30 min after oryzalin washout, which was strongly delayed and visible only after 60 min in GA-deficient (PAC-treated) roots (Figure 2A). We also analysed MT turnover rate using fluorescence recovery after photobleaching (FRAP). We found that the recovery of MT-associated TUA6-YFP was delayed in GA-deficient (PAC-treated) conditions as compared to GA-optimal (mock) conditions, despite the initial recovery rates being similar (Figures 2B and 2C).

**Fig. 2.**
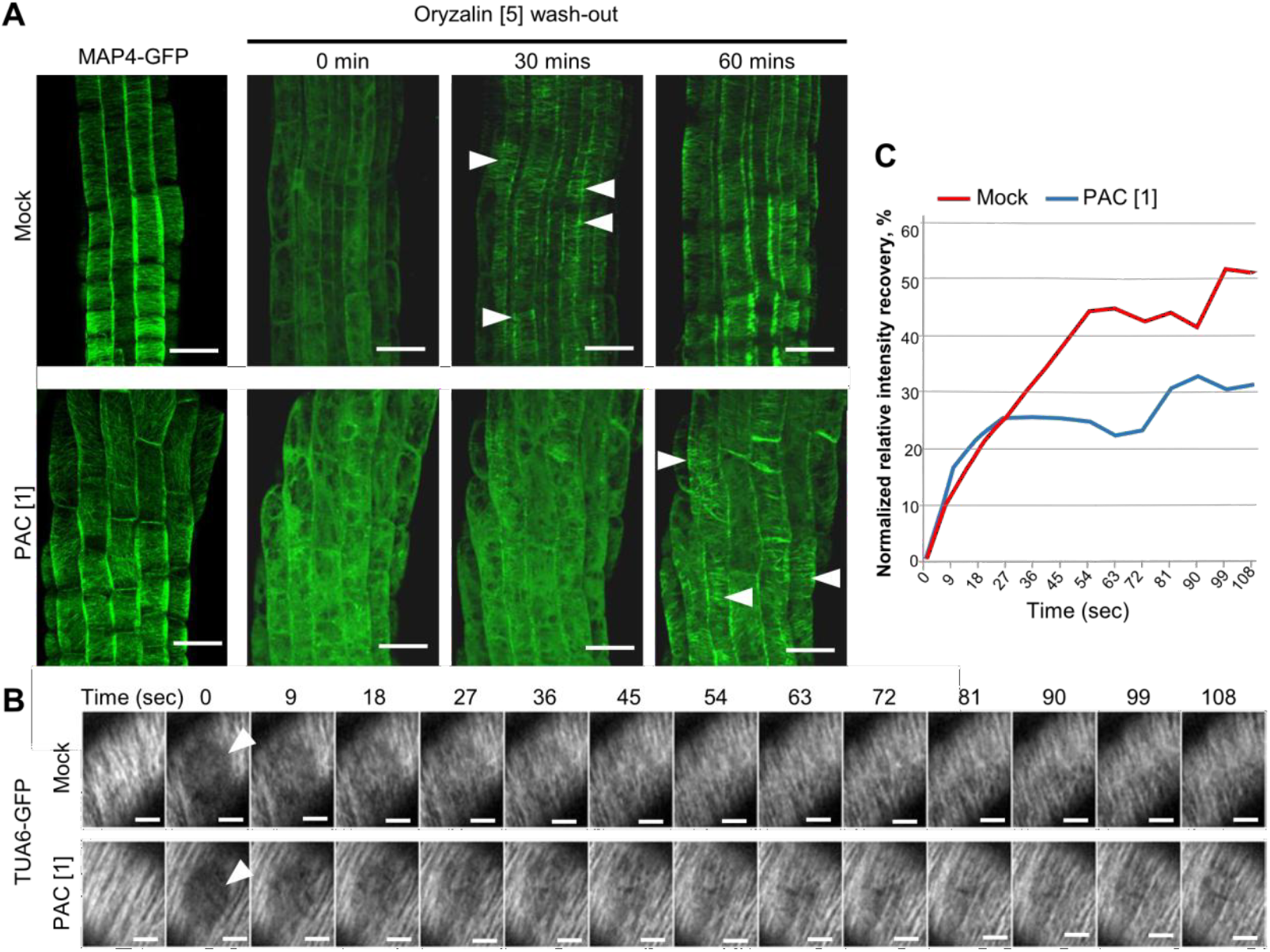
GA positively impacts microtubules turnover. (**A**) Cortical MT network in designated conditions as visualized in root epidermis by MAP4-GFP. The first visible appearance of CMTs structure is indicated by white arrowheads. (**B, C**) Fluorescence recovery after photobleaching (FRAP) in TUA6-YFP root epidermal cells. The photobleached area is indicated by arrowheads. Representative images (B) and quantification of MAP4 signal intensity (C). The intensities in the photobleached areas were measured in ImageJ. The t=0 was normalized as 0% and the recovery percentage was intensities on each time point relative to this value.

Together, these observations show that GA deficiency delays whereas GA supplementation promotes CMT recovery, suggesting a positive GA effect on MT network stability or tubulin mobility.

### GA promotes microtubule polymerization by a non-transcriptional mechanism

To further dissect the mechanism by which GA positively regulates MT network recovery or stability, we tracked MT plus end-binding protein EB1b (Chan et al., 2003) during GA-depleted and GA-supplemented conditions.

We performed time lapse visualization and of EB1b-GFP, and by using a max-projection of a small z-stack at each time point we were able to obtain EB1b-GFP “comets”, reflecting the growth rate of MT plus ends (Mathur et al., 2003; Schuyler & Pellman, 2001) (Figure 3C). The average velocity of MT polymerization was strongly reduced after 60 mins of PAC treatment from 0.16 microns/sec (GA-optimal) to 0.11 microns/sec (GA-deficient) (Figures 3A and 3D). In a complementary experiment, the application of GA on PAC-pretreated roots resulted in an increase of MT growth from 0.09 micron/sec to 0.17 micron/sec after 30 mins and reached 0.19 micron/sec after 60 mins of GA exposure (Figures 3B and 3D). The same EB1b-GFP time lapse imaging experiments in the presence of mannitol treatment, which blocks GA effect on cell expansion (see Figure 1C and 1D), revealed that the GA effect on MT polymerization occurs also when GA-promoted cell expansion was blocked by osmotic stress (Figure S1). These results show that GA depletion causes a decrease in the MT polymerization rate, while GA supplementation rapidly increases MT polymerization rate by a mechanism independent of cell expansion.

**Fig. 3.**
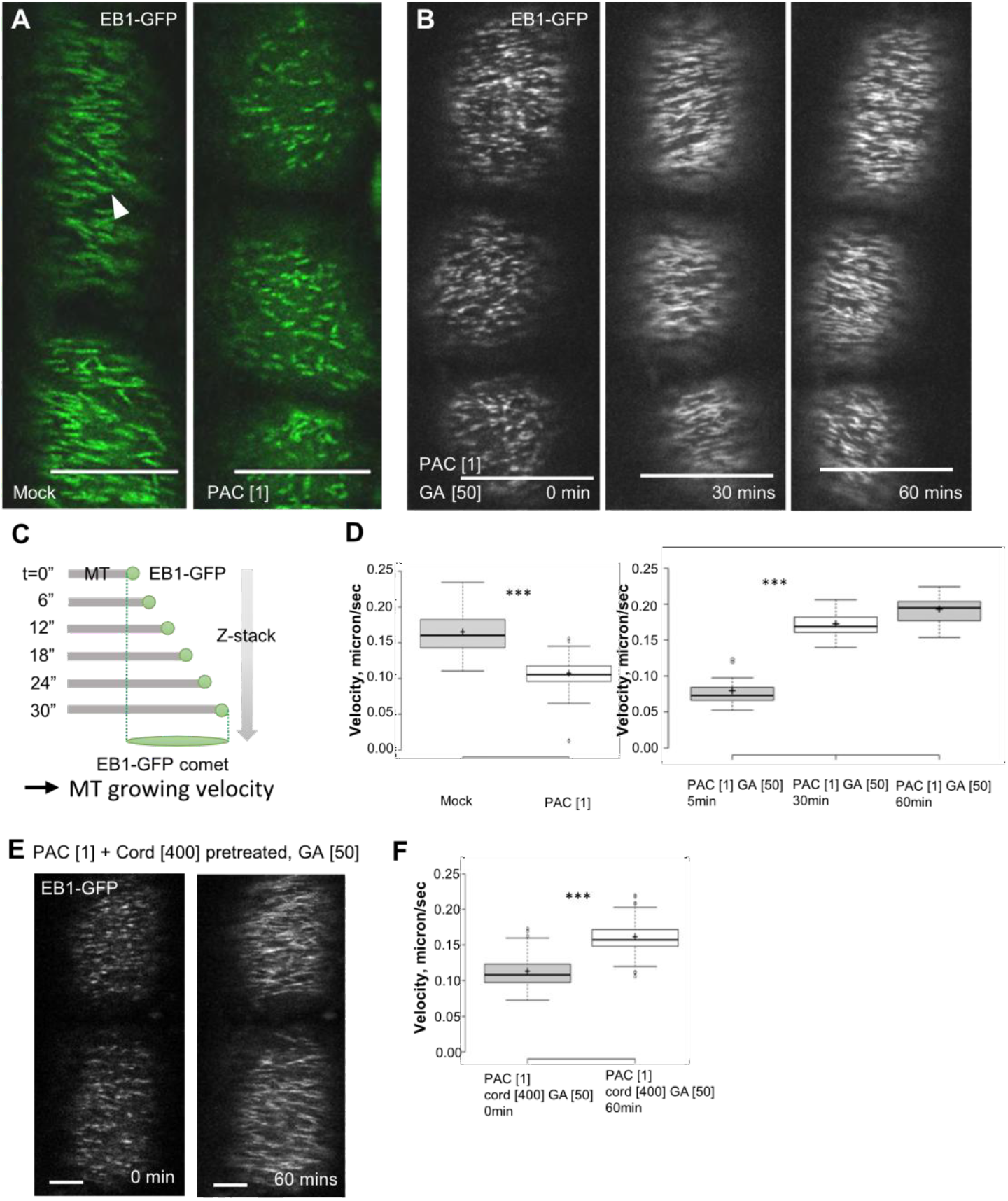
GA promotes microtubules polymerization by a non-transcriptional mechanism. (A-D) MT polymerization velocity in designated conditions as visualized in root epidermis by EB1b-GFP MT plus end marker. Images taken each 6 sec for 30 sec duration and projected by *Z*-stack maximum image projection to create comets used for the quantification of MT plus-end growing velocity (C). Representative images of the EB1b-GFP comets after GA depletion (1 μM PAC for 30 mins; A) or GA after addition (50 μM GA; B) with corresponding quantifications (D, E). (F, G) Transcription inhibitor cordycepin (400 μM) was added to the pretreatment medium with 1 μM PAC for 30 mins and the EB1b-GFP comets were tracked after supplementing 50 μM GA. Representative images (F) and quantifications (G). Statistics were performed using Mann-Whitney U-test and 2-tail student t-test. *** represents a significant difference (*p*<0.001). Scale bar = 10 μm.

The canonical GA signalling pathway across species and tissues converges on the nuclear DELLA transcriptional repressors and ultimately leads to transcriptional reprogramming that promotes various processes in plant development including cell growth (Cheng et al., 2004; Davière & Achard, 2013; Davière, De Lucas, & Prat, 2008; Dill, Jung, & Sun, 2001; Fu et al., 2002; Harberd et al., 1998; Richards et al., 2001; Sun, 2010; Wenzel & Wasteneys, 2000). Therefore, we tested whether the GA effect on MT polymerization occurs via transcriptional mechanism. We pretreated the roots simultaneously with PAC and inhibitors of transcription (cordycepin) (Siev, Weinberg, & Penman, 1969) or translation (cycloheximide) (Obrig et al., 1971) and then applied GA. Under these conditions, which inhibit transcription or translation (Li et al., 2021), GA was unable to promote cell expansion and root growth (Figure S2). This is consistent with GA regulating cell expansion by DELLA-mediated transcription (Cheng et al., 2004; Davière & Achard, 2013; Davière, De Lucas, & Prat, 2008; Dill, Jung, & Sun, 2001; Fu et al., 2002; Harberd et al., 1998; Mathur et al., 2003; Schuyler & Pellman, 2001; Wen & Chang, 2002) and also verifies that the inhibitors are effective.

Notably, under the same conditions, GA treatment still evidently accelerates the MT polymerization (from 0.11 micron/sec to 0.16 micron/sec after 60 mins) (Figures 3E and 3F). This further confirms that that GA effect on cell expansion and MT polymerization are independent and also implies that GA targets the MT polymerization by a non-transcriptional mechanism.

Overall, these observations revealed that GA, besides indirectly influencing MT arrangement through an effect on cell expansion, also directly targets MT polymerization by a cell expansion-independent, non-transcriptional mechanism.

### GA effect on microtubule polymerization involves DELLA degradation and prefoldins

Next, we addressed a downstream signalling mechanism, by which GA may regulate MT polymerization. The major GA signalling components are DELLA transcriptional repressors, which are degraded in response to GA treatment (De Lucas et al., 2008; Feng et al., 2008; Locascio, Blázquez, & Alabadí, 2013).

To test whether the GA regulation of the MT dynamics requires DELLAs, we crossed a *EB1b-GFP* line with a *GAI::gai-1-GR* line, which expresses a dominant negative, non-degradable version of the DELLA protein that is retained in the cytoplasm and transported to the nucleus only after treatment with the synthetic glucocorticoid dexamethasone (DEX) (Gallego-Bartolomé et al., 2011; Koorneef et al., 1985; Peng et al., 1997). DEX treatment in this line resulted in a reduction of MT growth from ∼0.20 micron/sec in non-treated control or DEX-treated *EB1b-GFP*/*WT* lines to ∼0.15 micron/sec (Figures 4B and 4D). This result not only shows that GA effect on MT polymerization is DELLA-dependent, but also implies that it is the nuclear DELLA presence required for this effect.

**Fig. 4.**
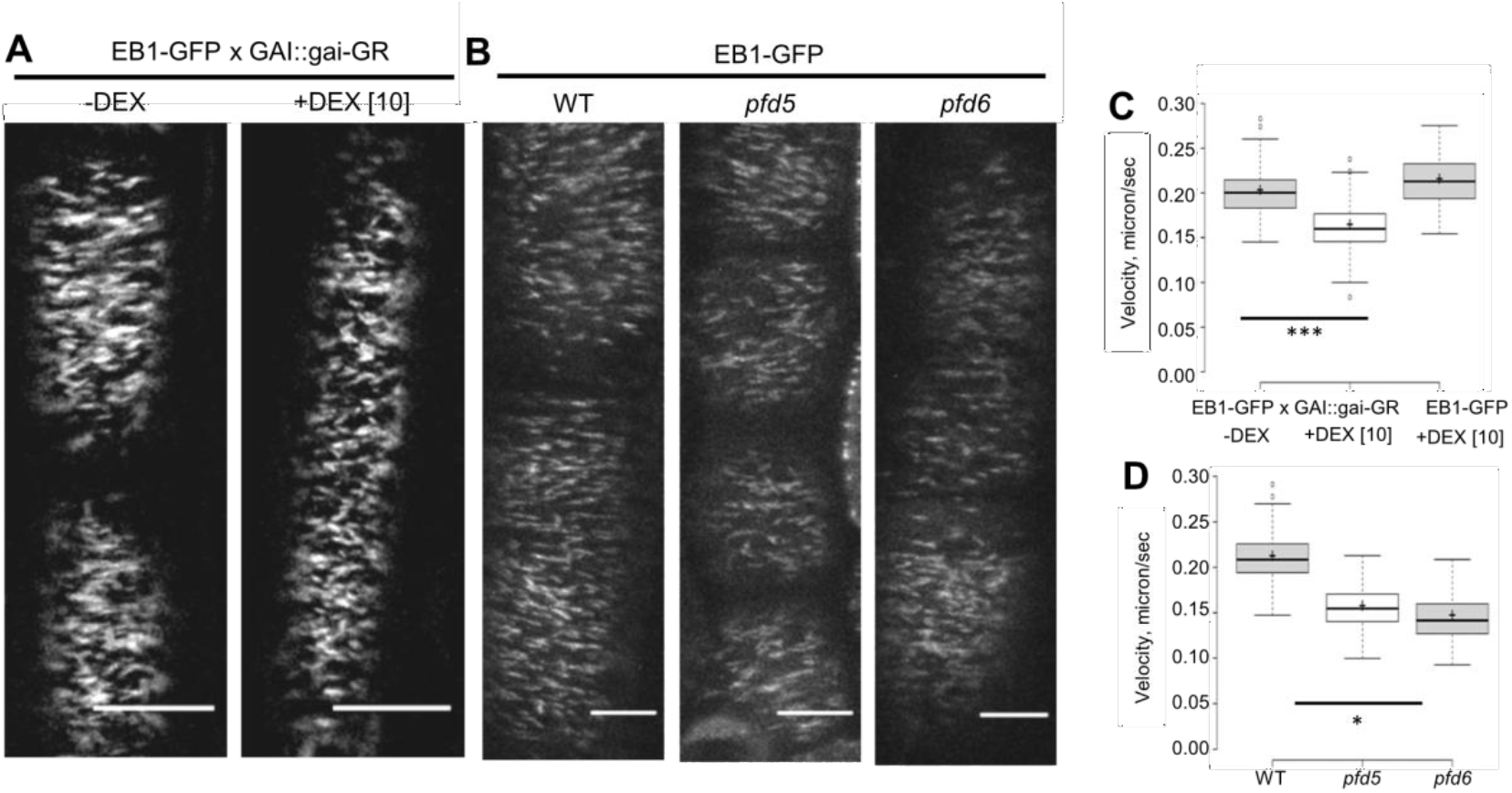
GA effect on microtubule polymerization involves DELLAs and prefoldins. (A) MT polymerization visualized as EB1b-GFP comet in *GAI::gai-GR* background in before and after induction with DEX, which induces a translocation of the stabilized, dominant *gai* DELLA protein from cytoplasm into nucleus. Scale bar = 10 μm. (B) MT polymerization visualized as EB1b-GFP comet in *pfd5* and *pfd6* mutants as compared to WT. Scale bar = 5 μm. (C, D) Quantifications of MT plus-end growing velocity in (A) and (B) from the length of EB1b-GFP comets as done in Fig. 3. Significant difference marked as *** representing *p*<0.001 in (C) and * representing *p*<0.05 in (D) was tested using a 2-tail student t-test.

Notably, DELLAs are also known to interact with and sequester in nucleus prefoldins (PFDs), which are important for tubulin folding and MT dynamics (Geissler, Siegers, & Schiebel, 1998; Locascio, Blázquez, & Alabadí, 2013; Vainberg et al., 1998). Therefore, we tested whether MT polymerization, as visualized by EB1b-GFP comets, requires PFDs by introducing the *EB1b-GFP* line into *pfd5* and *pfd6* mutants. We observed a strongly decreased MT growth (∼0.15 micron/sec) in both *pfd* mutants as compared to the WT plants, which showed a growth velocity of 0.21 micron/sec (Figures 4A and 4C). This shows PFDs’ importance for MT polymerization.

These observations show that the GA-induced enhancement of MT polymerization is mediated by DELLAs and their interaction partners PFDs. This suggests an existence of non-transcriptional DELLA-PFD-MT signalling module acting in parallel to the canonical, transcriptional DELLA-dependent signalling.

## Discussion

Plant growth requires that cell expansion and MT cytoskeletal organisation, which determines cell shape, remain tightly coupled. Our work reveals that phytohormone GA coordinates this coupling through two complementary mechanisms: a canonical transcriptional pathway that drives cell elongation, and a non-transcriptional mechanism that enhances MT polymerization. This GA duality underlies its role in growth enabling both, developmental stability and environmental responsiveness.

Our study demonstrates that GA-driven cell elongation is a cause of CMT reorientation. The early models hypothesize that MTs set the axis of growth (Shibaoka, 1994; Wenzel & Wasteneys, 2000) and it is evident that the well-established MT-dependence of cellulose synthesis (Lampugnani et al., 2018) determines the anisotropy of the cell wall (Majda et al., 2017) strongly influencing directional cell expansion. Our observations, however, show the opposite causality at least for a short-term effects: blocking elongation with mannitol prevented CMTs from reorienting, even when GA was present, whereas eliminating MTs with oryzalin or in *ktn1* mutant did not prevent GA from driving directional elongation. This is consistent with growth-induced changes in geometry and mechanical tension realigning MTs (Hamant et al., 2019; Hejnowicz, Rusin, & Rusin, 2000; Hoermayer et al., 2024) as shown also in case of auxin-induced hypocotyl growth (Adamowski, Li, & Friml, 2019). This interpretation reframes MTs not as upstream regulators of expansion but as sensors that emphasize growth direction once elongation is under way.

Beyond this growth-driven CMT reorientation, GA exerts a direct and rapid effect on MT dynamics. Manipulation of GA levels by GA addition or GA biosynthesis inhibition revealed that GA positively regulates MT dynamics as demonstrated by both faster recovery after photobleaching or after chemical-induced depolymerization. The live imaging of EB1b-GFP MT plus end marker (Chan et al., 2003) revealed that GA has a positive effect on MT polymerization on their plus ends. This happens regardless of the growth changes and does not require gene transcription or translation. Together this argues strongly for the existence of a non-transcriptional mechanism downstream of GA, which promotes MT polymerization.

Genetic analysis further revealed that this non-transcriptional effect requires canonical components of GA signalling, the DELLA transcriptional repressors, which are normally degraded in response to GA perception (Castro-Camba et al., 2022). Notably, it is the presence of stabilized DELLAs in the nucleus, which inhibits the MT polymerization. The other involved components are DELLA interactors, the Prefoldins, which act as cytosolic tubulin folding factors (Gu et al., 2008; Locascio, Blázquez, & Alabadí, 2013; Salanenka et al., 2018). Our observations thus confirm and extend previous notions that nuclear DELLAs can sequester Prefoldins by interacting with them and thus regulate MT dynamics (Locascio, Blázquez, & Alabadí, 2013).

This mechanism extends the role of DELLA beyond their classical function as transcriptional repressors (Davière & Achard, 2013; Dill, Jung, & Sun, 2001; Peng et al., 1997; Sun, 2010), positioning them as central integrators of cytoplasmic as well as nuclear responses. The DELLA-PFD axis thus illustrates how hormone signalling can non-transcriptionally regulate cytosolic processes including cytoskeletal arrays.

Taken together, these findings support a unifying model of GA signalling (Figure 5). The GA signals branches downstream of GA-induced DELLA degradation: (i) one branch derepresses transcription altering developmental programs for growth regulation; (ii) the other non-transcriptionally enhances MT polymerization via Prefoldins, ensuring rapid turnover of CMTs to align with growth and changed cell geometry.

**Fig. 5.**
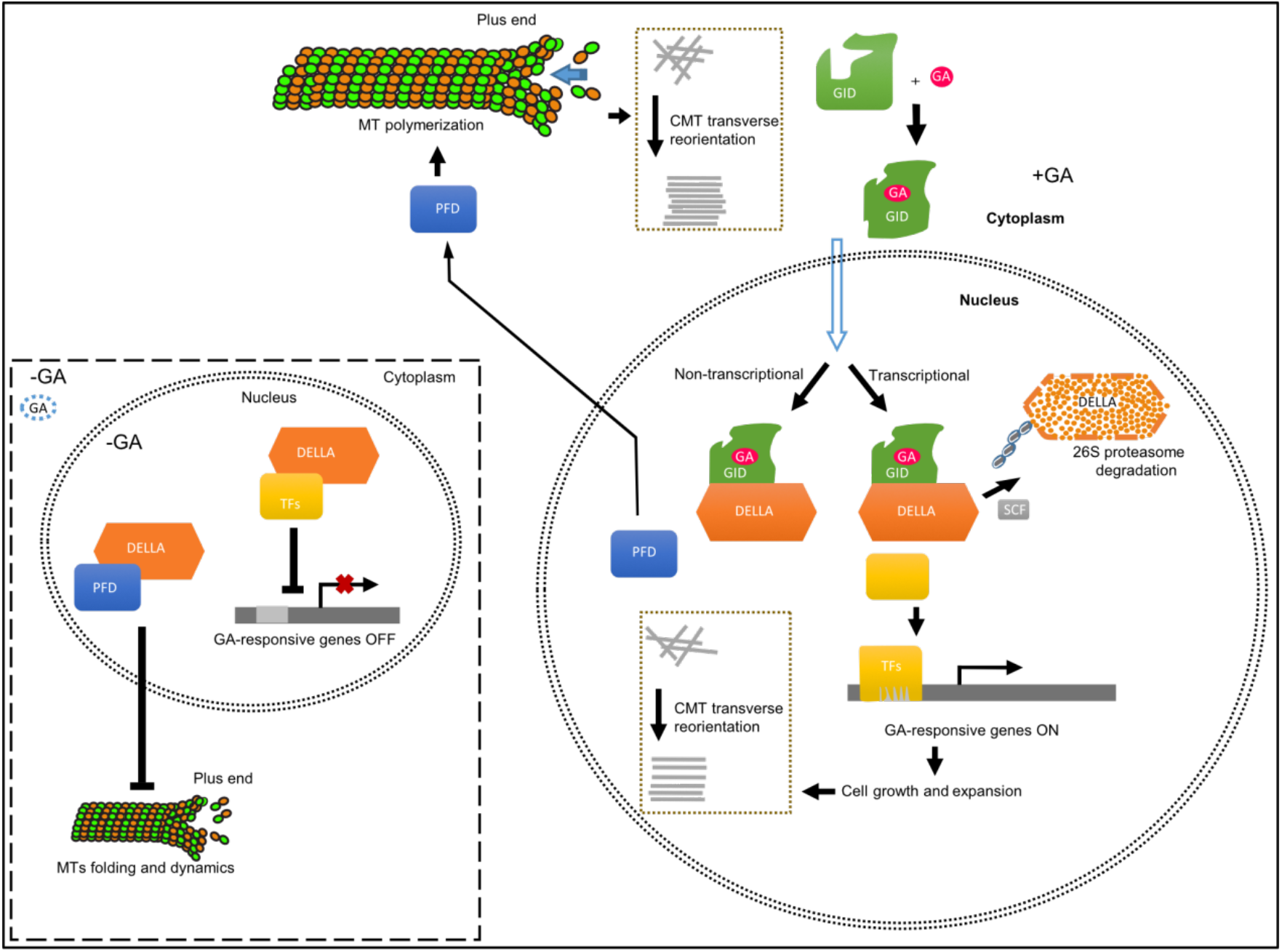
Mechanisms of transcriptional and non-transcriptional GA function. Proposed mechanisms how the GA signalling via GID-based perception mediates DELLA ubiquitination and degradation. This GA-induced DELLA degradation controls cell expansion and CMT reorientation via the canonical transcriptional mechanism. The parallel, non-transcriptional mechanism releases PFDs from the nuclear sequestration by DELLAs to promote MT polymerization in the cytosol.

Implications of these findings extend beyond Arabidopsis root growth. Such a dual signalling system represents a more general principle by which plants synchronize growth with cytoskeletal remodelling, enabling growth adaptation to fluctuating conditions, e.g. osmotic stress or mechanical disturbances, where plants must adjust growth direction without losing structural integrity. From an applied perspective, fine tuning this balance may allow us to promote growth without compromising mechanical strength, a trait of clear value in crops challenged by water deficits, wind or soil compaction (Achard et al., 2009; Hedden & Sponsel, 2015). Future work should dissect which exact PFD subunits and MT associated proteins act downstream of GA and how the non-transcriptional GA branch integrates with other hormones and signals that influence MT dynamics.

Our study extends the discussion of GA regulation of growth and MTs beyond correlative observations. We propose a framework where GA acts as a dual coordinator of growth and cytoskeletal organization. This integrative view underscores how a single hormone can unify long term development programs with immediate biophysical demands of the growing cell.

## Materials and Methods

### Plant material

*Arabidopsis thaliana* plants ecotype Col-0 were used in this work. The transgenic lines have been described before: *UBQ::TUA6-YFP* (Marc et al., 1998), *p35S::GFP-MAP4* (Schuyler & Pellman, 2001), *EB1b::EB1b-GFP* (Chan et al., 2003), *GAI::gai-GR* (Gallego-Bartolomé et al., 2011), *pfd5* (Rodríguez-Milla & Salinas, 2009), *pfd6* (Gu et al., 2008). The lines *GAI::gai-GR, EB1b::EB1b-GFP; pfd5, EB1b::EB1b-GFP*; *pfd6, EB1b::EB1b-GFP*; were generated by manual crosses and selected for homogeneous lines carrying both mutant allele and reporters.

### Growth Conditions

Seeds were sterilized overnight by 3% chlorine gas. Plants were grown vertically on half-strength MS medium (Murashige and Skoog, Duchefa) supplemented with 1% (w/v) sucrose and 1% (w/v) phytoagar (pH 5.7). Seedlings were grown on vertically oriented plates under a 16-h light/8-h dark photoperiod at 22/18 °C with LEDs-provided white light illumination. All experiments were carried out in the light period.

### Drug Application and Experimental Conditions

In the case of drug treatment, seedlings were moved to a MS plate or liquid MS medium containing the specified drugs for the specified duration. The final concentration of Paclobutrazol (PAC, Duchefa) was 1 μM, GA_3_ (Fluka) was 50 μM, mannitol was 300 μM, oryzalin was 5 μM, Cordycepin was 400 μM, and dexamethasone (DEX) was 10 μM. Mock treatments were done with equal amounts of solvent (DMSO/ethanol). A 30-min PAC-only or a pretreatment with the indicated drugs was followed by a concomitant drug as specified in the figures. All independent experiments were performed at least in triplicate on a minimum of eight individual plants.

### Imaging

Confocal images were taken with LSM 800 vertical-stage laser scanning confocal microscopes equipped with a 40× Plan-Apochromat water immersion objective. GFP markers were excited at 488 nm and emission was collected through band-pass filters 490–530 nm. For a vertical root tracking, 1× 3 tiles, 6 *z*-stacks (1 µm) were used for a 10-mins time-lapse recording. For quantifying the angle orientation of cortical MTs, a maximum intensity projection of confocal stacks was utilized to obtain images that were further processed and measured in the Zeiss Zen 2011 program, the ImageJ (National Institute of Health, http://rsb.info.nih.gov/ij), the Adobe Illustrator CC 2018, the GraphPad Prism 8 and the Microsoft PowerPoint programs.

### Quantification of microtubule orientation and plus end growth rate

FibrilTool plugin of ImageJ was used (Boudaoud et al., 2014) to quantify the cortical MT orientation in individual cells. Orientation was measured as an angle between cortical MT and the longitudinal root growth axis, with approximately 90º corresponding to the transversal orientation. For the MT plus-end protein (EB1b-GFP) tracking, an Andor spinning disk microscopy (CSU X-1, camera iXon 897 [back-thinned EMCCD], a 63× water immersion objective, FRAPPA unit and motorized piezo stage were used. Videos were acquired with a 500-ms exposure time, 5 *z*-stack 0.7 µm/each, every 6 sec, for 20-30 min.

EB1b-GFP spots were tracked by single-particle tracking (Tinevez et al., 2017) and the tracks were further analysed by using a script (Montesinos et al., 2020). To correct the sample drifting, the average motion direction of all spots at a given time point was calculated and smoothed over 10 frames. To quantify MT plus-end growth speeds, we compressed 5 z-stack images that were taken each 6 sec and obtained the EB1b-GFP comet as Figure 3C illustrated. The length of the GFP signals were measured manually and divided by duration to obtain the velocity.

### Quantification of cell expansion rate

For the quantification of cell elongation rate, roots of 5-day-old seedlings were monitored for 2 min, and 1 picture/30 sec was recorded as a single *z*-stack image with the LSM700/800 confocal microscope. The cell expansion rate was evaluated based on a relative increase in cell length that occurred within a 2-min time interval. For the cell expansion rate calculation, more than 10 cells per root and 3 to 6 roots per treatment in three independent replicates were collected.

## Supplementary Figures

**Suppl. Fig. 1.**
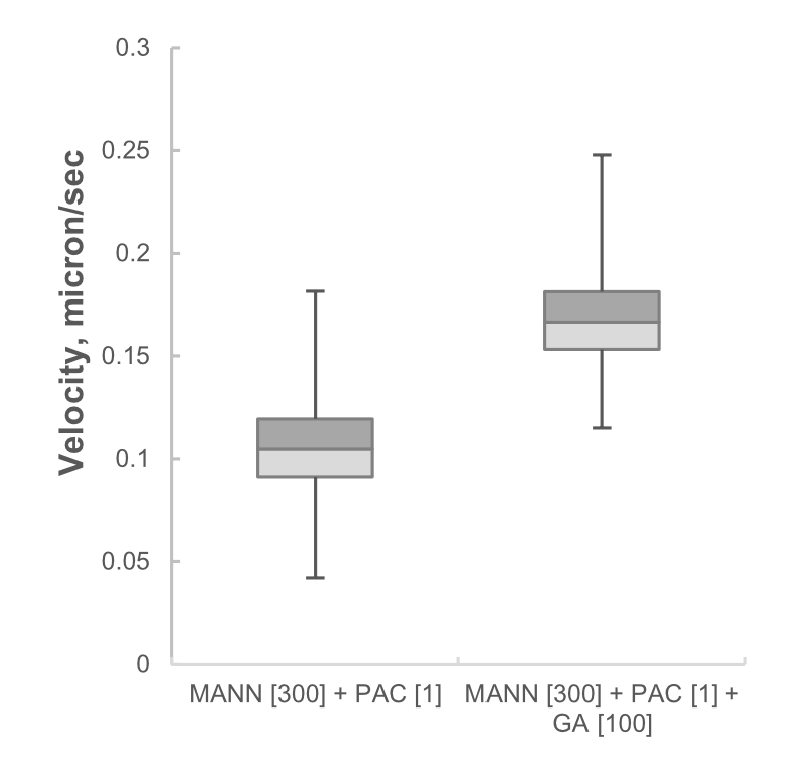
GA promotes cortical microtubules polymerization also in conditions of inhibited cell expansion. MT polymerization velocity in designated conditions as visualized in root epidermis by EB1b-GFP MT plus end marker comet quantified as in Fig. 3. Mannitol (300 μM) was added to the pretreatment medium with 1 μM PAC for 30 mins and the EB1b-GFP comets were tracked after supplementing 100 μM GA.

**Suppl. Fig. 2.**
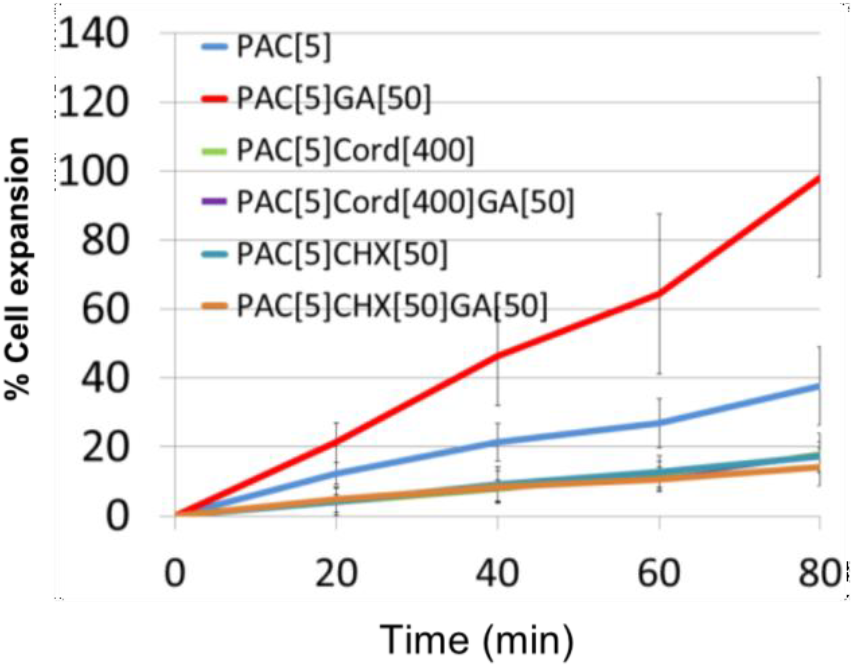
GA-induced cell expansion requires intact transcription and translation. GA effect on cell expansion in epidermis of root elongation zone. Seedlings were pre-treated with 5 μM PAC in presence or absence of transcriptional inhibitor cordycepin (400 μM) or translational inhibitor cycloheximide (50 μM) for 30 mins followed by 50 μM GA treatment as indicated. The percentage of cell expansion in all observed elongated epidermal cells was calculated by dividing the cell length by its original cell length.

